# Building Back More Equitable STEM Education: Teach Science by Engaging Students in Doing Science

**DOI:** 10.1101/2021.06.01.446616

**Authors:** Sarah C R Elgin, Shan Hays, Vida Mingo, Christopher D Shaffer, Jason Williams

**Affiliations:** Department of Biology; Washington University in St. Louis; St. Louis MO; Natural and Environmental Sciences Department; Western Colorado University; Gunnison, CO; Department of Biology; Columbia College; Columbia, SC; DNA Learning Center; Cold Spring Harbor Laboratory; Cold Spring Harbor NY

## Abstract

The COVID-19 pandemic is a national tragedy, one that has focused our attention on both the need to improve science education and the need to confront systemic racism in our country. We know that active learning strategies, in particular research experiences, can engage and empower STEM undergraduates, effectively closing the achievement gap for historically excluded persons. The apprenticeship model for STEM training – supervised research under a dedicated mentor – is highly effective, but out of reach for most students. Recent efforts have demonstrated that Course-based Undergraduate Research Experiences (CUREs) can be an effective approach for making STEM research accessible for all. Our meta-analysis of CUREs finds that published examples now cover the breadth of the typical undergraduate biology curriculum. A thoughtfully designed CURE can go beyond foundational knowledge and analytical thinking to include career-related skills, *e.g*., teamwork and communication. Similarly, it can be designed with equity as a foundational principle, taking into account the unique contributions of all students and their varying needs. We provide here an example framework (The “*Do Science Framework*”) for making STEM training more effective and inclusive using CUREs. While CUREs do not inherently address equity, there can be no equity in STEM education without equal access to research participation, and progress toward this goal can be achieved using CUREs. However, implementing new CUREs is not a trivial undertaking, particularly at schools with high teaching loads and little or no research infrastructure, including many community colleges. We therefore propose a National Center for Science Engagement to support this transition, building on experiences of current nationally established CUREs as well as the work of many individual faculty. In the aftermath of the COVID-19 pandemic, academia has a renewed responsibility to dismantle structural inequities in education; engaging all STEM students in research can be a key step.

## Introduction

### The immediate need for better science education

The Soviet launch of Sputnik in 1957 shocked the US public, precipitating a crisis in the country’s confidence in its education system. Congress promptly took action, passing the 1958 National Defense Education Act (US Senate Record). Coupled with increased funding for the National Science Foundation (NSF), the combined support for students and teachers generated a successful nationwide investment in science education (Mazuzan, 1994).

The COVID-19 pandemic is a national tragedy. The pandemic has raised anew national questions about our ability to effectively lead in science, a critical issue for national prosperity and security. It has also compelled us to confront the systemic racism that has led to disproportionate death and economic damage in Black, LatinX and indigenous communities, and has devastated an already deeply inequitable education system. Pandemics are not the only threats we face. Climate change, cyber-security, and other looming threats place renewed urgency on growing both the STEM workforce and a scientifically literate citizenry (Tilghman *et al*., 2021). To prevail, we need to revisit lessons learned from Sputnik and confront truths revealed by COVID-19. Sputnik showed that when the public is engaged, we can rise to the challenge. However, post-Sputnik education reforms benefited some groups and not others (Malcom-Piqueux, 2020). We need to avoid repeating that mistake and use this moment of national focus to dramatically improve science education while dismantling structural and systemic inequity and racism (**Box 1**).

#### Box 1: This image represents the difference between equity and equality

Equity is the fair treatment of each individual based on their needs and requirements, while equality is treating everyone the same regardless of needs (Espinoza, 2007). In the equity image, the different boxes represent the support provided based on the resources needed for the student to engage with the content (fruit). To achieve equity, the level or type of support (depicted by the boxes the students are standing on) is not evenly distributed to all students but provided according to need (Paul, 2019). One observation illuminating the failures of the current system is that students from historically excluded populations are more likely to change their majors and not complete a STEM degree; forty percent of Black students and twenty-nine percent of LatinX students drop out even though they enter STEM at the same rate as their white counterparts (Riegle-Crumb *et al*., 2019). **We have chosen to use the term historically excluded persons to represent the underrepresentation of LatinX, Black, and Indigenous people of color (BIPOC) in STEM fields.** One of the goals of the proposed National Center for Science Engagement is to remedy this numerical underrepresentation by building CUREs with equity as a foundational tenet in design and implementation. Image Credit: Saskatoon Health Regions, 2014.

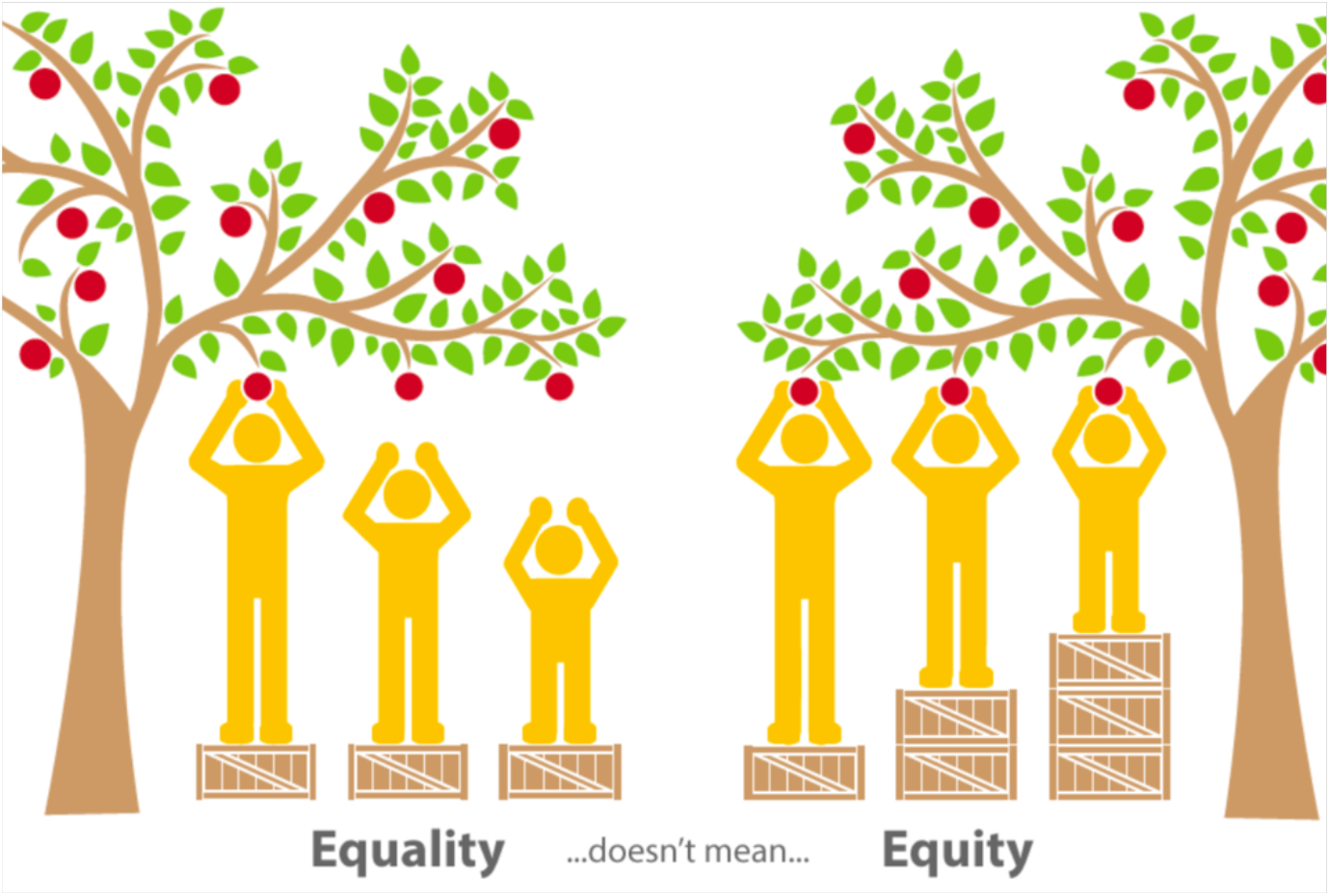

### We know strategies that work

The U.S. has developed resources to address these challenges. Over the last decade, the science and education communities together have identified evidence-based learning strategies that support *all* students in becoming engaged and successful in STEM. The recommendations in “*Vision and Change*” (AAAS, 2011) provide a framework for bringing these active learning strategies into the classroom. Those recommendations have already spurred significant innovation in science education (AAAS, 2019; NASEM, 2021). Studies show that engaging undergraduates with active learning experiences is effective, increasing their involvement and agency in their own science education (Ballen *et al*., 2017). These strategies also help close the performance gap for historically excluded students (Theobald *et al*., 2020). Thus, we propose basing key reforms on the central concept of ***teaching science by engaging students in doing science.*** We refer to this as the ***Do Science Framework***.

### A focus on undergraduate education

While we argue that this strategy is critically important to K-16 for developing a scientifically literate citizenry, it is even more essential for developing the larger and more diverse science and engineering workforce that we need. As delineated in the PCAST 2012 report, improving undergraduate STEM education to retain a larger fraction of the students who enter college/university with STEM interests would be a major step forward that we can accomplish. Traditionally the *lab apprenticeship* has been the gateway to a career in science. Mentored undergraduate research experiences improve retention of students in STEM majors, improve graduation rates (National Research Council, 2003; Laursen *et al*., 2010; AAAS, 2011), and increase interest in pursuing a STEM PhD (Seymour *et al*., 2004; Russell *et al*., 2007; Villarejo *et al*., 2008; Jones *et al.*, 2010; Espinosa, 2011; Hernandez *et al.*, 2013). Such programs, found in both research universities and colleges, including HBCUs, are equally effective for historically excluded students (Hrabowski *et al.*, 2011; DiBartolo *et al*., 2017; Katz *et al*., 2017). Unfortunately, the traditional structure of STEM apprenticeships often makes them inaccessible and unwelcoming for many historically excluded and non-traditional students (Bangera & Brownell, 2014). Further, by their nature these programs can only serve a small population of students, and often focus on those students who have already exhibited interest and enjoyed some success in STEM in high school. Apprenticeships simply cannot scale to meet the current need: there are not enough mentors, lab space, or summer fellowships available to provide for all STEM undergraduates, resulting in a system of selection that generally lacks equity. Fortunately, a wide range of alternatives are being explored (NASEM, 2017).

### A CURE: Course-based Undergraduate Research Experiences

A major strategy for reaching large numbers of students is to implement Course-based Undergraduate Research Experiences (CUREs). PCAST 2012 developed the importance of morphing lab courses into research experiences, creating CUREs. CUREs (or CREs) are typically one- to three-semester courses where student teams explore a scientific question through a research project (NASEM, 2015; Elgin *et al*., 2016; Dolan, 2016; Dolan & Weaver, 2021). (**Box 2**; also, see the Supplement for descriptions of many successful CUREs.) CUREs have been found to increase retention and graduation in a STEM major, having a similar impact for historically excluded students as for the whole student population (Rodenbusch *et al*., 2016), A direct comparison of the impact of research experiences through CUREs and through an apprenticeship found similar affective gains for students using either path, while indicating that participation in CUREs can reduce the achievement gap between high-performing students and their peers (Shapiro *et al*., 2015).

#### Box 2: Defining a CURE

A CURE should involve students in 1) use of scientific practices, 2) discovery, 3) broadly relevant or important work, 4) collaboration, and 5) iteration (multiple rounds of problem solving and troubleshooting). “Scientific practices” include asking questions, building and evaluating models, proposing hypotheses, designing studies, selecting methods, using the tools of science, gathering and analyzing data, identifying meaningful variation, navigating the messiness of real-world data, developing and critiquing interpretations, and communicating findings (Auchincloss *et al*., 2017; Linn, 2015; CUREnet). While CUREs vary considerably in practice, all include most of the above features in some way.

The following are suggested as characteristics of a good project to be developed as a large-scale *Do Science CURE*, starting from the discussion in Hatfull *et al*., 2006:

- Sub-dividable into “parallel projects” so that faculty can teach common methods, with students carrying out individual projects (*e.g*., each 2-student team isolating their own phage), generating student ownership; students can report their findings as a poster or brief paper, such as a *microPublication* (see Raciti *et al*., 2018 for latter). Individual projects are assembled into a whole for meta-analysis and joint publication;
- Technically simple – to provide low costs, safety, high probability of success, and fit with scheduling constraints – or central provision of any hi-tech element;
- Conceptually accessible – varies with targeted student level;
- Providing multiple achievement outcomes (*e.g*., student skill gains, incremental results obtained) so that students build confidence over the semester;
- Amenable to a quality control system to support student findings (*e.g*., two independent determinations and reconciliation of any differences);
- A goal of reporting findings to a wider audience, whether scientific or local community.

**We believe that now is the time for a national campaign to facilitate adoption of CUREs at *<u>all</u>* undergraduate institutions, and that this effort must be grounded in a meaningful framework for equity.**

Implementing CUREs broadly does not by itself do all that is needed to advance equity in science education; but we argue that there can be no equity without access for all students to research experience, making this an essential first step. The ***Do Science Framework*** we propose is centered on engaging all STEM undergraduates in basic and/or applied research as a significant part of their academic-year curriculum. Such a transformation of undergraduate STEM education can help address educational, economic, and health inequities and contribute to sustainable national prosperity and security.

## Methods

### Literature review

Given current crises in human health, food security, and the environment, we focus here on the life sciences, but similar arguments apply to STEM as a whole. To assess the feasibility of this proposal, we have undertaken a systematic review of the literature, including consulting major projects such as CUREnet.

Using elements 2-9,17, 23a-d, 24b, and 25-26 of the PRISMA checklist (Page, 2020) we report on a qualitative review of available literature on CUREs. We had two research questions: 1) which biology subdisciplines in the typical undergraduate biology curriculum are served by CUREs (as defined in Box 2); and 2) which CUREs can be readily expanded to engage a large number of students. The search protocol started with all authors developing a consensus list of subject areas to represent the breadth of undergraduate biology. From November 2020 – March 2021, all authors searched the current most complete resources on CUREs (*e.g*., CUREnet), as well as the major journals for reporting on undergraduate biology education (*e.g*., CourseSource, CBE Life Sciences Education), and common search databases (SCOPUS). In addition, we circulated these results to a group of 15 faculty experts in undergraduate biology research experiences, requesting their review for significant oversights or omissions.

We generated a list of 54 published and communicated CURES using this methodology. For the examples in Table 1 we selected CUREs that provided sufficient description to implement the CURE (*e.g*., the curriculum, experimental protocols, etc.). Many CURE publications focused solely on the description of an assessment of that CURE and so were excluded. For the examples in Table 2, we selected documented CUREs that are operating at a national level and/or can be scaled up locally to engage large numbers of students.

**Table 1.** CUREs for (nearly) every biology course. Based on literature review, we present selected examples of scalable CUREs that cover the most common subject matter of the undergraduate biology curriculum.

| Course | Research Subject | Lab Techniques (reference) |
| --- | --- | --- |
| Introductory biology | Local species of choice | Ecological sampling, molecular biology, DNA barcoding (Hyman *et al.*, 2019) |
| Introductory biology | Phage discovery | Ecological sampling, microbiology, molecular biology, bioinformatics (Hanauer *et al.*, 2017; Jordan *et al.*, 2014) |
| Aquatic ecology | Aquatic food web | Stable isotope analysis (Carroll *et al.*, 2019) |
| Bioinformatics | Gene annotation | Use of evidence tracks in a genome browser; BLAST (Lopatto *et al.*, 2008; Shaffer *et al.*, 2010; Shaffer *et al.*, 2014) |
| Cell biology | Ovarian follicle cells | Dissection and microscopy (Tootle *et al.*, 2019) |
| Conservation biology | Urban diversity of birds | *In silico* analysis of online data from citizen scientists (Gastreich, 2020) |
| Developmental biology | Zebrafish embryogenesis | Bioinformatics, zebrafish maintenance, RNA extraction, RT-PCR (Felzien 2016) |
| Ecology | Species dispersal | Field work, DNA barcoding, phylogenetic analysis (Elwess *et al.*, 2018) |
| Genetics | Mutational analysis | Site-directed mutagenesis, screening (Walsh, 2020) |
| Microbiology | Soil microbe diversity | Field work, molecular biology, bioinformatics (Lo & Mel 2017; Shapiro *et al.*, 2015) |
| Molecular biology | Gene expression | RT-PCR, *in situ* hybridization, phylogenetic analysis (Shapiro *et al.*, 2015) |
| Neurobiology | *C. elegans* neurons | RNAi screen, behavioral analysis (Kowalski *et al.*, 2016) |
| Plant ecology | Plant-bacteria interactions | Plant growth studies, bioinformatics, gene annotation (Shapiro *et al.*, 2015) |

**Table 2.** Examples of CUREs that can scale to large numbers of students. For those interested in a particular project, websites are indicated; all websites checked as of April 15, 2021. See Supplemental Materials for more information.

| Name | Topics | Website | Location |
| --- | --- | --- | --- |
| DNA Barcoding | Ecological sampling, DNA barcoding, conservation genomics, biodiversity | dnabarcoding101.org | National; Cold Spring Harbor Laboratory DNA Learning Center |
| San Diego Biodiversity Project | Ecological sampling and DNA barcoding | sdbiodiversity.ucsd.edu/info | UC San Diego |
| SEA-PHAGES | Microbiology and genome annotation | seaphages.org | National; University of Pittsburgh / HHMI |
| Genomics Education Partnership | Bioinformatics and genomics | thegep.org | National; University of Alabama |
| Competency-based Research Laboratory | Virology, development, or microbiology | crlc.ucla.edu | UCLA |
| BASIL Biochemistry Consortium | Protein biochemistry | basilbiochem.github.io/basil | National; RIT New York |
| Freshman Research Initiative | Various | cns.utexas.edu/fri | UT Austin |
| First-year Research Immersion | Various | binghamton.edu/first-year-research-immersion | Binghamton University |
| The First-Year Innovation & Research Experience | Various | fire.umd.edu | University of Maryland |
| Prevalence of Antibiotic Resistance in the Environment | Microbiology and molecular biology | sites.tufts.edu/ctse/pare/ | Tufts University |
| SIRIUS Project | Stream ecology | csus.edu/college/natural-sciences-mathematics/sirius/ | CSU Sacramento |
| Tiny Earth | Microbiology | tinyearth.wisc.edu/ | National; University of Wisconsin |
| Small World Initiative | Microbiology | smallworldinitiative.org/ | National; New York, 501(c)(3) |
| Vertically Integrated Projects | Various | vip.gatech.edu | Georgia Institute of Technology |
| Vertically Integrated Projects | Various | engineering.purdue.edu/VIP | Purdue University |

## Results

### Findings

We find that CUREs have been developed for every area of biology commonly taught (**Table 1**). Examples in Table 1 are taken from the published literature. The breadth of topics expands if one also considers CUREs reported as part of a larger programs (see Table 2, freshman initiatives), or on CUREnet and in other less formal collections. We also find that there are many good examples of CUREs that have been scaled up to engage large numbers of students, either on a given campus or in a national consortium; examples that are well described are given in **Table 2**. Topics related to genetics and ecology are particularly well represented, perhaps because those methodologies can often benefit from a large, scientifically trained workforce. In addition, research projects in these areas are readily designed to address topics of concern to students in human health, climate change, etc.

## Discussion

### Enabling CUREs at all colleges/universities

Despite the growing evidence of the effectiveness of CUREs, and their utilization at all types of institutions, including community colleges (Hewlett, 2018), they do not yet appear to be widespread (AAAS, 2019). Designing and implementing an effective CURE is not a trivial undertaking, particularly for faculty at institutions with high teaching loads and low research resources. A number of organizations have emerged to assist faculty, including several “national CUREs” (see Table 2 and **Box 3**). Such projects have been shown to enable faculty across all types of institutions to incorporate research into their teaching (Lopatto *et al*, 2014). But many more such projects are needed to achieve national scale. Therefore, we propose creation of a **National Center for Science Engagement (NCSE)** to stimulate and assist in the development and support of both existing and new national CUREs, as well as supporting and to support the efforts of individual faculty to transform their lab course into a CURE (*e.g*., Dolan & Weaver, 2021). Such a Center will serve equity goals by reducing the burdens of major curriculum reform, support that is particularly important for resource-limited schools. An NCSE should provide not only access to various projects (with supportive curriculum, faculty training, etc.) but also additional centralized resources and guidance aimed at reducing and eliminating inequity. We invite the scientific community to consider what science, relevant to the field and/or the public, might be accomplished with the help of “massively parallel undergraduates,” and the mutual benefits that can result.

#### Box 3: Real-world examples of CUREs that can scale

The SEA-PHAGEs program, specifically for freshmen (Jordan *et al*., 2014; Hanauer *et al*., 2017; Staub *et al*., 2016), the Genomics Education Partnership (GEP) (Shaffer *et al*., 2010, 2014; Elgin *et al*., 2017), the Tiny Earth project (Hurley *et al*., 2021), DNA barcoding projects (Marizzi *et al*., 2018; Hyman *et al*., 2019), and others (see the Supplement) are designed to scale to multiple institutions and can engage large numbers of students. These projects have identified *a central question* where gathering multiple samples across the country, or otherwise pooling the efforts of large numbers of undergraduates, makes possible science that could not otherwise be done. A central website provides the scientific rationale, lab protocols, critical tools for data collection and/or analysis, sample curriculum, and group communication tools. The central organization provides start-up faculty training and continuing peer support, and often plays a key role in publication of the pooled results. This allows faculty to step in and join the effort fairly quickly (Lopatto *et al*., 2014). Contrasting mechanisms useful for many campuses include GCAT-SEEK (Genome Consortium for Active Teaching Using Next-Generation Sequencing) (Buonaccorsi *et al*., 2014) and DNA Barcoding developed by the Cold Spring Harbor Laboratory DNA Learning Center, which provide a central source of tools and peer support but have each faculty member develop their own project using these tools; and Vertically Integrated Projects (VIP) (Coyle *et al*., 2014, 2015), which provides organizational templates and support, but relies on each campus to devise its own research projects based on faculty interests. Many such projects engage students long-term, sophomore through senior year. All of these examples provide the individual faculty member with a “community of practice” (Lave & Wenger, 1991) to help make the transition to teaching a CURE. The SEA-PHAGEs program, which stresses building project ownership, scientific community values, development of science identity, and scientific networking uses the term “iREC,” inclusive Research Education Community, to describe themselves (Hanauer *et al*., 2017).

### Capturing the Potential: The “*Do Science*” Framework

The *Do Science Framework* draws on current practice in CUREs (PCAST, 2012; NASEM 2015, 2017; Dolan & Weaver 2021) and expands to include direct attention to equity, workforce development, and connection to community. We identify four elements of this framework:

#### Promote comprehensive research projects organized around a variety of topics

Projects should meet the definition of a CURE given in Box 2.

1. **Equity: start with freshmen, then follow up**. Start all students with a core freshman research experience designed to be interesting and accessible to potential science majors and non-science majors alike. The pedagogical goals of this course are to introduce students to research and to illustrate the breadth of biology. Students should leave knowing that they are capable of doing scientific research (Killpack *et al*, 2020; Cooper *et al*, 2020). This freshman introduction should be followed by opportunities to participate in topic-oriented CUREs, vertically integrated projects (VIPs), or other research experiences, internships, etc. every semester thereafter (typically one course each semester for a science major). See **Boxes 4** and **5.**
2. **Workforce development: Go beyond foundational knowledge**. Broaden curricula beyond the foundational knowledge required in the domain science. Include digital literacy and crossdisciplinary knowledge as needed for the project, while providing experiences in science communication, teamwork, and other “soft skills” that employers value (NASEM, 2018). Also include elements of humanistic knowledge (responsibility to ethics, equity and societal concerns) (Mishra *et al*., 2020).
3. **Connection to community: Utilize both basic and applied research**. *Do Science* projects should increase student engagement both by including topics of immediate interest and by revealing links between basic research and problems of broad community interest. A CURE can be based on community partnerships, addressing a local ecological problem or local business/industry need (see examples among VIP teams at vip.gatech.edu/teams). Many CUREs centered on basic research questions may not have an apparent connection to an immediate problem that a student may want to solve; however, these connections can usually be made with faculty guidance as students realize how basic science precedes applied science, for example in phage research (Dedrick *et al*., 2019).
4. **Equity: Involve everyone**! Previous science education reforms have focused more on course content and teaching methodology than on the larger and systemic problems of inequity. As NSF has long recognized, scientific merit is intertwined with broader impacts. We assert that educational reforms must now center inclusion and equity (Asai 2020). A challenge we must address as a nation is how to achieve this centering. We anticipate this will take a variety of forms, including culturally relevant curriculum design, new models for resource allocation, as well as institutional and professional development.

##### Box 4: How would the *Do Science* concept work for beginning students?

Valentina is enrolled in her first science course at the regional community college with the goal of becoming a registered nurse. She is immediately introduced to challenging scientific problems. The college freshman biology course is organized around a research experience using DNA barcoding to explore the local ecosystem. To get ideas, Valentina and her teammates visit the National Center for Science Engagement website where they review examples of projects other students have done. They become interested in research on fire ants, which are a new problem in their state. She spends the first several weeks working collaboratively with her team to learn the lab techniques and acquire background on the investigation they will join, contributing data to a national research effort to develop a fire ant species range map. The bulk of the semester is spent in a mix of reading scientific literature, designing/carrying out experiments, collecting data, and communicating her research group’s results. While her time on campus and in the field is limited, all of the resources and much of the work are accessible online, allowing her to participate fully while still balancing family needs. Although the work does not always go smoothly, the online TAs are helpful, and the guiding faculty member is supportive. Time is allotted to reconsider, redesign and repeat data collection, experiments, etc. as needed. She is exposed to many areas of biology she never knew existed, including bioinformatics and genomics. While her whole team wishes they could have done more, they are satisfied that they made progress on the problem and learned much more about biology than they perhaps expected. Valentia is excited to share what she has learned with her family members, particularly her cousins on the farm, and is thinking that the effort to pursue a science-based career may be worth it. She also learns that the genomics technology she learned about is increasingly important in medicine. Valentina is now thinking about expanding her career interests, and looks into the NCSE’s website, which has information on job opportunities in genomics-based medical diagnostics. She is looking forward to the sophomore genetics course, which centers on cancer genetics/genomics and offers a research project in Drosophila genomics.

##### Box 5: How would the *Do Science* concept work for advanced students?

After his freshman experience, Ahanu debates his choices – biotech vs environmental work, semester-long CUREs vs a VIP-style team. He chooses to join a team working on water quality and ultimately graduates from his four-year Tribal College with a degree in environmental science. He starts out a bit behind his peers but, by sticking with one project over several years, he is able to develop a more comprehensive understanding of the relevant science and is asked to help teach new members of the team. While he cannot answer all the questions the beginners have, he can answer many of them, and when he cannot he asks the faculty member in charge and learns more about the project. He builds confidence in his own abilities and gains newfound leadership skills as he works with the intro students. Ahanu can remember his grandparents talking about the uranium mining that occurred on the Navajo Nation reservation during the 1940s (see also Middlecamp, 2004). He vividly remembers his parents’ frustration with the lack of safe drinking water for their community. The chance to investigate the impact of mining on water quality, an issue relevant to him and his community, is appealing. Ahanu enjoys being able to apply what he is learning through his research to a cause that he is passionate about and enjoys being a part of the VIP-style team for several years. With faculty mentorship, he is able to “do science”and integrate it with local history, health equity and environmental justice issues. Students in the course are engaged in reading the literature while carrying out their research on best practices for monitoring water quality and treating contaminants; they also have the opportunity to attend talks by activists from the Native American Water Justice organization. Ahanu plans to present his work at the American Indian Science and Engineering Society Regional and National Meetings. He hopes to continue this work in graduate school and eventually plans to work for a tech company, improving detection and mitigation of contaminants in water.

The *Do Science Framework* strives to move CUREs to the next stage by placing them at the center of a reorienting of the undergraduate STEM major and by focusing attention on equity.

### Supporting *Do Science*: A Proposal for a National Center for Science Engagement

#### Equality of access as the path towards equity

CUREs are the natural successor to the apprenticeship model because they can scale (see Box 3). To overcome the barriers to implementing CUREs and create a new science education landscape that can work for all students, we propose a new infrastructure to support undergraduate STEM education: a **National Center for Science Engagement (NCSE).** The National Center’s primary goal will be to provide support, as detailed below, for implementing CUREs at all two-year and four-year US colleges and universities. This should include the development of CUREs at all levels – from facilitating the ongoing work of current large multi-site collaborative CUREs and cultivating the development of new national CUREs on diverse topics, to helping innovative faculty that seek to create new CUREs tuned to local expertise and local student interests. Supportive frameworks that facilitate development of campus-specific efforts, whether VIP multi-semester research teams or one-semester CUREs built around individual faculty research interests, should be included. Thus, the NCSE will sponsor a growing portfolio of national CUREs that a faculty member/school can join, while also supporting local efforts, in part by providing the centralized services listed below.

#### Functions of a National Center for Science Engagement

Taking a nationwide approach to undergraduate STEM education will require the NCSE to provide several services:

1. Efficiently curate a portfolio of national CUREs open to all faculty, supporting those CUREs through the dissemination of “best practices” and implementation strategies in alignment with the *Do Science* framework. Centrally coordinating CURE design should lower the barrier for developing new CUREs by sharing lessons learned, rather than expecting scientists with a good idea for a national CURE to solve every challenge on their own. New national-scale CUREs might engage students in contributing data to develop a national profile, as in the bird counts organized by the Cornell Lab of Ornithology (birds.cornell.edu/home/citizen-science-be-part-of-something-bigger) (Gastreich, 2020), or might engage students in analyzing large data sets, as in genome annotation (Elgin *et al*, 2017); there are many possibilities. The NCSE should encourage development of CUREs that address national needs for data collection (*e.g*., ecosystem monitoring) and/or workforce development and national security (*e.g*., developing familiarity with AI).
2. Provide professional development for faculty including science background (*e.g*., training on new methodologies and technologies pertinent to a new CURE),) and equity (*e.g*., training on incorporating ethics and anti-racism in curricula). The latter training can work in concert with mandates many institutions are developing to ensure inclusive classrooms, presenting solutions to challenges in the context of STEM teaching (Ero-Tolliver, 2019; Walsh *et al*., 2013; Killpack & Melón, 2016).
3. Coordinate access to cyberinfrastructure, working with projects such as QUBES, Galaxy, and CyVerse to provide workshops that provide faculty with the needed data management skills to meet the technical needs for the research of interest. This training should address the significant barriers faculty face in this area, particularly; in biology for example this would include guiding faculty new to bioinformatics (Williams *et al*., 2019).
4. Adopt or develop appropriate tools for program evaluation, and provide faculty training in educational assessment, as appropriate (Auchincloss *et al*., 2014).
5. Facilitate student-to-student interactions. While providing a research experience, national CUREs also have the benefit of potentially connecting students across institutional and geographic boundaries (see Boxes 3-6).
6. Develop public outreach that connects research experiences to communities and develops public interest and engagement, including connections to K-12 enrichment and community/citizen science programs such as the Citizen Science Association (citizenscience.org).

It is critical that the NCSE maintain an equity-first approach by supporting the *Do Science* concept and related science education practices at institutions that face high barriers to adoption. This could be accomplished in several ways:

1. Leadership should be driven by an advisory group that includes stakeholders from primarily undergraduate institutions (PUIs), community colleges, Historically Black Colleges and Universities (HBCUs), and Hispanic-Serving and Tribal Colleges. Care should be taken to also address the needs of historically excluded students enrolled in predominantly white institutions.
2. Projects in the CURE portfolio managed by the Center, while grounded in the science, should connect directly or indirectly to problems that are important to the communities, and those connections should be made visible to the students. For example, the NCSE could maintain a “bulletin board” for “problems that we need solved,” helping to recruit scientists interested in developing a CURE to address those problems.
3. The Center should drive institutional change, buy-in and adoption of CUREs at schools new to this format by including faculty training as above, access to low-barrier national projects, and perhaps access to implementation funding (microgrants).

#### The need for a National Center to support the *Do Science* framework

While CUREs are broadly effective, the ability to implement this strategy is largely dependent on the resources available at a given institution, leading to the perpetuation of structural inequality (**Figure 1, A**). Further, how students proceed after a research experience is again solely dependent on institutional resources. Some CUREs (B) are part of a national network (*e.g*., SEA-PHAGES, GEP, DNA Barcoding); these provide professional development opportunities, allowing faculty to engage with a community of practice. CUREs operating in this framework have been able to successfully broaden participation by developing accessible research projects and actively recruiting faculty at under-resourced institutions. The durability and reach of these projects are largely dependent on funding to maintain the central organization, with many projects disappearing after the end of a grant. (C) A National Center for Science Engagement would provide a coordinated and enduring structure for CUREs, equitizing access, and providing support for institutions and students along the lines described above.

**Figure 1:**
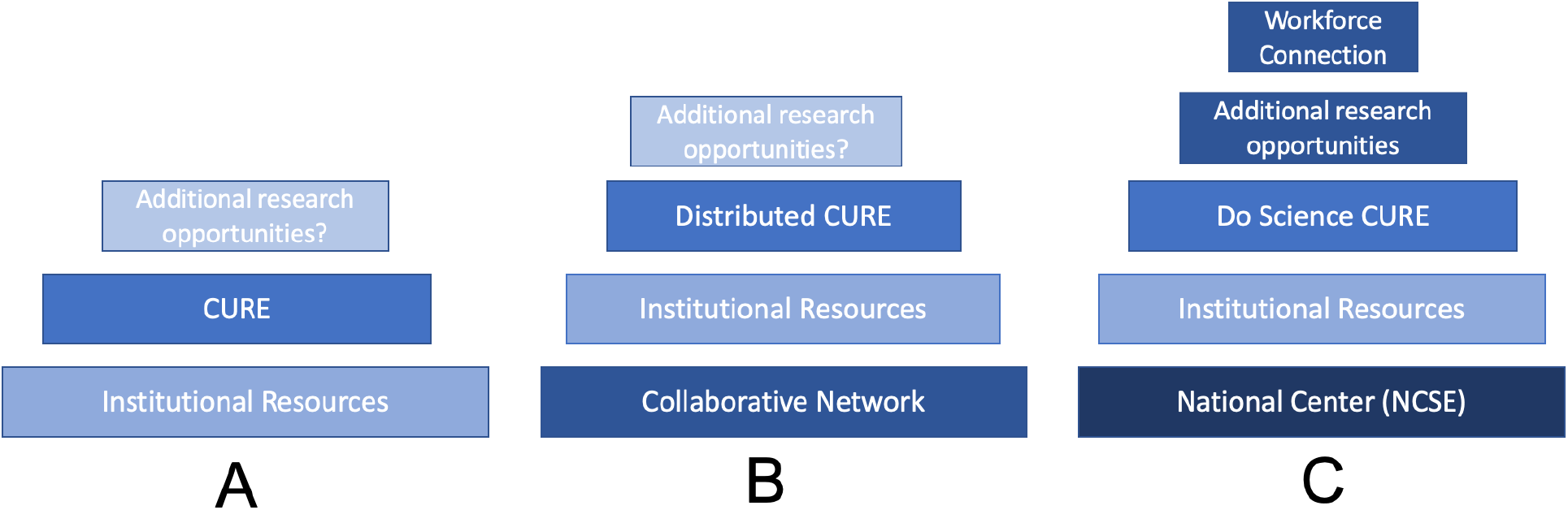
Placing CUREs in the *Do Science* framework. CUREs are broadly effective (A) but can gain strength from being part of a national partnership (B); the available resources for faculty/institutions could be strengthened through a National Center for Science Engagement (C).

#### Integrating *Do Science* CUREs with the National Center for Science Engagement

Rethinking how we do Freshman Biology and supporting that effort with a National Center for Science Engagement, could transform our current efforts. The NCSE can remove burdens from faculty and institutions in a costeffective manner while developing the additional layers needed to promote research experiences that enrich students and the community. In **Box 3** we document several examples of CUREs that can be readily extended into the *Do Science* framework. Projects based on analysis of data available online have the further advantages of low laboratory costs (requiring only computers and internet connectivity), no lab safety issues; and open access 24/7; are often amenable to peer instruction/coaching; and allow for mistakes that are inexpensive in dollars and time, allowing so students have more freedom to learn (Elgin *et al*., 2017; Lopatto *et al*., 2020). There are masses of data available free online (for example sequenced genomes; ecosystem monitoring data; satellite survey data) begging for analysis, and exploitation of these resources to address questions of interest could be developed. National CURE designs need to be such that students do not face enrollment barriers such as high lab fees or uncertainty as to how they earn the letter grade they would like to receive (NASEM, 2015). The best approaches are those equally accessible to large state campuses (thousands of undergraduates) and to community colleges, ensuring successful articulation (NASEM, 2017). Providing systematic support for CUREs through the NCSE will have benefits for students, faculty, institutions, and society (**Box 6**).

##### Box 6: Benefits from instituting *Do Science*

###### Benefits for students

- Increases availability and reduces barriers to gaining a research experience;
- Creates opportunities to gain practical skills and experience in active problem solving, critical thinking, quantitative analysis and communication;
- Provides students entrée into a “community of practice” and all the attendant benefits of the social capital gained by membership;
- Provides a way to make a meaningful contribution with lasting value.

The above benefits increase the chances that the student will graduate with a STEM major (Rodenbusch *et al*.,2016) which translates into higher estimated future earnings (Walcott *et al*., 2018).

###### Benefits for faculty

- Provides a means to combine teaching interests and responsibilities with research efforts and opportunities for authorship (Shortlidge *et al*, 2016);
- Creates a baked-in pipeline of well-trained students to bring into individual research labs (post-CURE participation);
- Creates access opportunities to join a “community of practice” STEM peer group in an existing field of research, with concomitant access to ongoing training and up-to-date, validated curriculum in a fast-moving field (Lopatto *et al*., 2014);
- Provides escape from the “tyranny of content,” allowing students to meaningfully engage with the biological concepts underlying the research investigation, maintaining their natural curiosity (Petersen *et al*., 2020).

###### Benefits for administrators and colleges/universities

- Builds a culture of faculty-student interactions;
- Increases faculty participation in scholarly activities;
- Can create opportunities to partner with local businesses and community groups;
- Drives greater retention overall (Rodenbusch *et al*., 2016), maintaining the tuition stream.

###### Benefits for society

- Addresses racial and social inequities in STEM education, with a focus on MSI and rural institutions, as CUREs can increase student engagement/motivation/achievement and have been shown to close gaps between white and BIPOC students (Rodenbusch *et al*., 2016);
- Addresses the workforce needs for a large, diverse pool of STEM trained individuals sufficient to meet the demands of a flourishing economy (PCAST, 2012; Tilghman *et al*., 2021); proactively addresses, in a national and comprehensive manner, the continued underrepresentation of BIPOC in the STEM workforce (Corneille *et al*., 2020; Coleman, 2020);
- Can address local environmental concerns, support local businesses, as well as basic research of national significance;
- Accomplishes important research that could not be done in any other way by leveraging the work of thousands of undergraduates working together with their mentors to collect and/or analyze large amounts of data (*e.g*., Pope *et al*. 2015; Dedrick *et al* 2019; Leung *et al*. 2015, 2017). One can readily visualize projects that will help assess the impacts and improve mitigation of global warming, provide environmental data leading to better management of local watersheds, etc.

#### Organizing the National Center

NSF and other funding agencies could generate appropriate calls for proposals to identify and support potential *Do Science* CUREs within the NCSE, in some cases working up to a level that meets a national demand, while in other cases ensuring equitable access for a smaller but equally meaningful audience. For example, CUREs developed with NSF IUSE funding might naturally evolve to become part of the NCSE. Different agencies, including foundations, government agencies, and industry groups, might wish to support national CUREs in particular areas, both to achieve mission goals and to support work-force development. Support for a pedagogically successful project should be renewable until the scientific and educational goals are accomplished. A competitive proposal will engage students in a “good project” as described above; will support faculty professional development for implementation, including mentorship and equity; will provide needed resources and curriculum; and will have a plan for development of a community of practice. The NCSE’s resource allocation mandate would help institutions that serve historically excluded persons center the science that matters to their communities. In short, prototypes for the *Do Science CURE* exist and they work; this mechanism now needs to be made equitable, scaled up, and institutionalized at a national level using a “collective impact” approach. NSF INCLUDES might provide a starting model (includesnetwork.org/home).

## A Call for a Convening

According to *A Brief History of NSF* (Mazuzan, 1994), “Even before the Soviets put Sputnik I in orbit on October 5, 1957, [NSF] and American scientists had been concerned with the state of American science vis-a-vis the Soviet Union. While the satellite provided the first human reach beyond the planet, it symbolized in America the need for improving scientific education and basic research, needs already known to the scientific community.” Recent workshops on the future of STEM demonstrate that many are again concerned about that future; see for example *The Future Substance of STEM Education*, fall 2020 (Mishra P, 2020); and *Symposium on Imagining the Future of Undergraduate STEM Education*, November 12-13, 2021 (NASEM 2021).

With calls for change to our educational system building for decades, every agency that funds science and education, every faculty member, and every member of the public needs to ask the question -– if not now, when? For science education in the US, COVID-19 surpasses the precedent of Sputnik. The STEM education community needs to recapture the momentum that propelled the United States then, as well as fulfill an obligation to channel it in a way that dismantles inequity.

Recognizing that a real solution will require significant community input, we are calling for a convening to explore and refine the solutions suggested here and elsewhere, as well as ideas that have not yet been heard. We propose a convocation on **Building Back Undergraduate STEM: Using CUREs in an Equity-first Approach to Doing Science**. This convening could be formatted into four meetings:

1. A gathering of interested stakeholders (faculty, research scientists, funders, etc.) to frame the issues described above, in particular examining approaches to promote equal access to research opportunities and equal achievement for all students; reviewing the criteria for good projects; determining whether a coordination center (The National Center for Science Engagement) is warranted, and if so, the desired scope for the Center;
2. A gathering of research scientists, engineers and policy experts, partnered with representatives from business/industry and local/state/national governments, to propose CUREs that the National Center might sponsor, addressing national challenges in basic and applied science that fit the descriptions addressed above;
3. A gathering of a diverse group of faculty and students, including leaders from community colleges, PUIs (large and small), HBCUs, and Tribal colleges, to determine the kinds of intellectual and physical support that should be part of an NCSE-supported CURE; and to refine foundational guidelines for equity, examining how research experiences can best serve the needs of diverse students;
4. A synthesis meeting of participants from the above sessions with potential funders to discuss pilot projects and the programmatic infrastructure needed to implement a decadal vision.

We invite the scientific community to consider what science, relevant to the field and/or the public, might be accomplished through the efforts of “massively parallel undergraduates” that cannot be accomplished in any other way. Careful community input will be needed to ensure that historically excluded voices are heard throughout the process, from the start in setting the agenda to the final synthesis. We invite the community’s feedback, input, and direction in taking the next steps. If you are a faculty member, representative of a current CURE project, or other stakeholder who supports the broad vision proposed here and would like to assist in its development, please comment here and/or contact the corresponding author. We anticipate that the next step will be to seek funding for the convocation described above.

## Supplemental Materials

### Item 1: Descriptions of CUREs with the potential to scale to large numbers

#### BASIL Biochemistry Consortium

The BASIL (Biochemistry Authentic Scientific Inquiry Laboratory) biochemistry consortium provides the infrastructure and curriculum to allow students to analyze proteins with known structure but unknown function. BASIL’s computational analyses and wet-lab techniques are designed to be used by students in biology, biochemistry, chemistry, or related majors and are often used as part of upper-level laboratory coursework with at least one semester of biochemistry as a pre-requisite or co-requisite. However, the curriculum is flexible and can be adapted to other appropriate courses and to match the available facilities, the strengths of the instructor, and the learning goals of a course and institution.

#### Genomics Education Partnership

The Genomics Education Partnership (GEP) provides training, resources, and mentorship to its members, allowing them to provide courses that include research projects in genomics and bioinformatics for their students. The GEP partners with researchers whose projects require that a group of genes or genomic regions be carefully annotated. Students develop their annotation and describe supporting evidence based on their analyses of evidence tracks in a genome browser that is maintained by the GEP. After reconciliation of multiple annotations from different students, the final annotations are used by the science partners to carry out the planned meta-analysis/investigation. Current projects are looking at the evolution of the insulin signaling pathway using 28 species of Drosophila; evolution of the venom proteins in parasitoid wasps; and the impact of genome expansion of the F element (a heterochromatic chromosome arm) on genes and genome architecture. Students can present their individual work in a poster at a meeting or in a *microPublication*; GEP faculty and contributing students are eligible to be co-authors on the publications that result from meta-analysis utilizing their contributions.

#### DNA Barcoding

Similar to the universal product code (UPC) that identifies each consumer product, a “DNA barcode” is a unique pattern of DNA sequence (~700 nucleotides in length) that can potentially identify each living thing. In a typical experiment, DNA is isolated from a specimen, a barcode region is amplified, and the DNA sequenced. Then the barcode sequence is used to search for its closest match in GenBank, the international DNA database. DNA barcoding may be particularly well suited to freshman CUREs, as it integrates big ideas from molecular biology, genetics, bioinformatics, ecology, and biodiversity – while at the same time providing the flexibility to address a variety of student-driven questions. Barcoding can be mastered in a relatively short time, allowing students to generate new data and reach a satisfying research endpoint within a single course. Furthermore, many freshmen have limited patience for bioinformatics, so DNA barcoding provides a wet-lab or field-based “hook” to increase engagement. In 2011, the Cold Spring Harbor DNA Learning Center developed an integrated biochemical and bioinformatics workflow for student DNA barcoding including low-cost reagents, kits, protocols that eliminate most specialized equipment, and a classroom-friendly web-based bioinformatics platform (DNA Subway). Based on user registrations, we estimate that 12,000 college students annually conduct DNA barcoding projects using DNA Subway. Remarkably, James Madison University has adapted the DNALC’s barcoding curriculum to reach over 1,700 freshman students per year in semester-long research (Hyman 2019). DNA barcoding and metabarcoding have been used as an end-to-end research program with 2,900 high school students from 176 schools in the New York metro area, reaching high numbers of historically excluded students. To date, these students have identified food fraud, tracked invasive species, conducted bio-inventories of local parks and published 160 DNA barcodes for species not previously represented in GenBank (Marizzi *et al*., 2018).

#### SEA-PHAGES

This two-semester project begins with students collecting a soil sample to find new viruses, first using classical microbiology techniques to plaque-purify their phage, then characterizing that phage by noting growth characteristics, generating a restriction map, and (where possible) obtaining an EM picture. A growing database allows them to make a tentative classification. Phage are sequenced over the winter break, and in the spring, students learn to use a variety of bioinformatics tools to annotate their phage. Students closely identify with “their” phage - and have the opportunity to name it. Not only do the students report their results in undergraduate research symposia, the fully annotated phage can be submitted for publication; the acquisition of large libraries of phage genomes enables comparative and evolutionary studies which make a significant contribution to the scientific literature. The impact on students is very positive (*e.g*., Jordan *et al*., 2014). The course emphasizes principles of microbiology and bioinformatics but depending on the instructor’s design can incorporate basic principles of ecology, molecular biology, cell biology, biochemistry, enzymology, structural biology, bioinformatics, and evolution. While the requirement for wet-bench work (and sterile plates) may seem limiting, economy approaches have been developed by the practitioners, and this course has been used to serve half of the freshman class at Pitt (G. Hatfull, personal communication).

#### Small World Initiative

The Small World Initiative provides the instructional material and instructor training to allow faculty to guide students in isolating bacteria from soil in their local environments for further laboratory experimentation. By testing their bacteria against clinically-relevant microorganisms, and characterizing those that show inhibitory activity, students’ research has the potential to uncover novel antibiotics.

#### Tiny Earth

Tiny Earth provides resources to allow instructors to guide students in “student-sourcing antibiotic discovery” from the soil. Students carry out fieldwork to isolate new strains of microorganisms, which they subject to genomic and metabolomic studies in the lab in order to discover novel bioactive-producing microorganisms. Promising new strains are submitted to the Tiny Earth Chemistry Hub, where they can be subjected to further chemical analysis. Students can also develop and test their own hypotheses regarding variables that they think might influence antibiotic production in bacteria. The results from this research are often presented within the schools where the work takes place. The program also consists of an educational component to raise awareness about antibiotic resistance and the appropriate use of antibiotics.

#### Freshman Research Programs

Different schools have developed different versions, but all are based on using a team approach to involve students in research right at the beginning of their college careers, providing a smorgasbord of topics to choose from. Involvement generally lasts from one to three semesters. In some cases, students apply for or are invited to participate in these programs, so these opportunities are not *per se* available to all, while in other cases the format is more open. Regardless, the programs provide the participating students with an opportunity to engage in a faculty-mentored research project early in their college/university career.

#### Vertically Integrated Projects

An alternative to building CUREs around current courses is to build CUREs around the research interests of the faculty, either in conjunction with a course they teach, or as a separate course, potentially organized as a Vertically Integrated Project (VIP). In the VIP format, undergraduates join long-term project teams led by faculty working in their area of scholarship and exploration. Students can join a team in their sophomore year and engage continuously through their senior year, experiencing different roles, from beginner to student expert. This system can reduce the burden on the faculty member, because senior students mentor the incoming students. While long-term immersion in a single project reduces the breadth of the student’s education, it does provide a setting in which the student can make substantial contributions. This format also lends itself to interdisciplinary projects.

